# AlphaScreen profiling of extracellular vesicle-associated platelet, immune, and vascular markers reveals disease-specific signatures in ME/CFS, fibromyalgia, and long COVID

**DOI:** 10.64898/2026.11.10.765688

**Authors:** Maria Ljungström, Esther Masiá, María Jesús Vicent, Elisa Oltra

## Abstract

**Background/Aim:** Myalgic encephalomyelitis/chronic fatigue syndrome (ME/CFS), fibromyalgia (FM), and long COVID (LC-19) are chronic diseases with overlapping clinical features and poorly understood pathophysiology. Increasing evidence suggests the involvement of immune dysregulation, endothelial dysfunction, and altered vascular physiology. Extracellular vesicles (EVs) are key mediators of intercellular communication and are envisioned as sensors of homeostasis and disease. The aim of this study was to determine whether EV marker profiling could provide a basis for the differential diagnosis of ME/CFS, FM and LC-19 and/or patient stratification.

**Methods:** EV-enriched plasma from ME/CFS, FM, cases co-diagnosed with ME/CFS and FM, LC-19 patients, and matched healthy controls (n = 10 per study group) were analyzed. EV concentration and size distribution were assessed by nanoparticle tracking analysis, followed by targeted profiling of EV-associated surface markers via AlphaScreen, including tetraspanins CD9 and CD63, platelet-(CD41/CD61), myeloid-(CD13), immune-(CD74 and CX3CR1), and renin-angiotensin system-associated ACE1 (also known as CD143), and ACE2 surface markers.

**Results:** The studied disease groups presented distinct EV signatures. LC-19 samples exhibited the most pronounced and coordinated alterations, characterized by increased levels of CD9/CD74, CD9/CX3CR1, and CD9/ACE1-positive EVs, along with reduced overall particle concentrations. Notably, platelet-derived CD9/CD41/CD61-positive EVs were significantly elevated in both LC-19 and ME/CFS, as compared to healthy controls, indicating enhanced platelet activation. Immune-associated markers revealed differential pathway activation in LC-19, with CD74-positive EVs reflecting antigen presentation and inflammatory signaling, and CX3CR1-positive EVs indicating altered leukocyte trafficking and vascular interaction. Increased ACE1-positive EVs in LC-19 further suggested a shift toward pro-inflammatory renin-angiotensin signaling.

**Conclusion:** Collectively, these findings support a disease-specific dysregulation of platelet, immune, and vascular pathways. The integrated EV signatures distinguish LC-19 from the related chronic conditions ME/CFS and FM, highlighting the potential of EV-based profiling for differential diagnosis. Our results provide a systems-level perspective on the differential pathophysiology of these disorders and identify EV-associated markers as candidate biomarkers with clinical value.

**Highlight:** Extracellular vesicle profiling identifies disease-specific signatures in ME/CFS, fibromyalgia, and long COVID, enabling highly accurate discrimination of long COVID from clinically overlapping conditions (AUC = 0.914).

**GRAPHICAL ABSTRACT:** 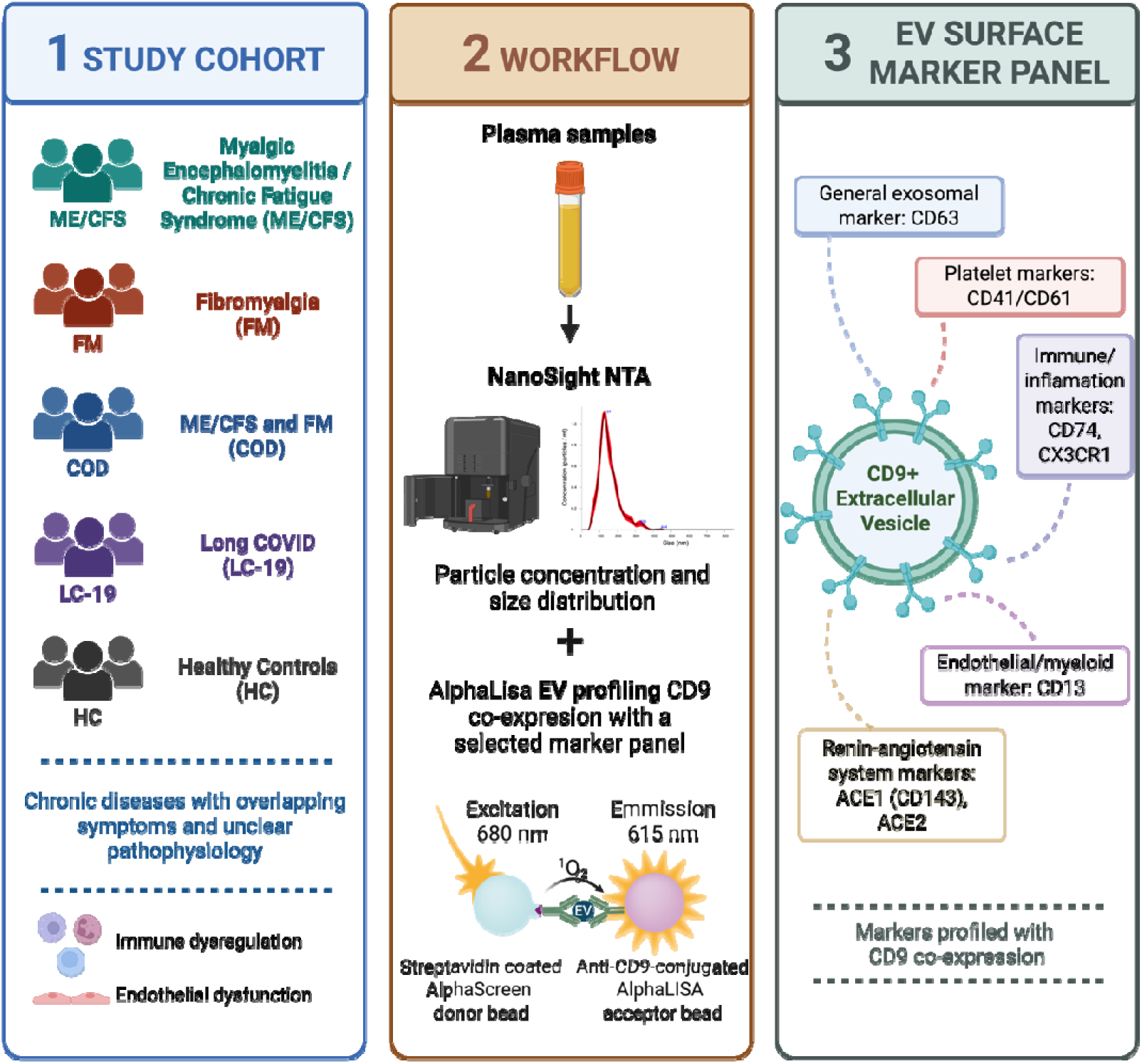

## INTRODUCTION

Myalgic encephalomyelitis/chronic fatigue syndrome (ME/CFS) (ICD-11 8E49), fibromyalgia (FM) (ICD-11 MG30.01), and long COVID (LC-19) (ICD-11 RA02) are complex chronic conditions characterized by multisystemic symptoms, including fatigue, pain, and cognitive impairment (1–6). Despite their prevalence and their relevant clinical burden, the underlying pathophysiology of these conditions remains poorly understood, currently lacking objective biomarkers for their diagnosis. The presence of shared clinical features and potential comorbidities further complicates diagnostic uncertainty, highlighting the need for approaches that capture underlying biological differences.

Increasing evidence suggests that these diseases involve dysregulation of multiple interconnected systems, including immune activation, endothelial dysfunction, and altered vascular homeostasis.

Chronic inflammation, immune cell activation, and microvascular alterations have been proposed as contributing factors in ME/CFS, FM and the co-diagnosis of both (COD), while LC-19 has been associated with persistent immune dysregulation, endothelial injury, and thrombo-inflammatory processes following SARS-CoV-2 infection (5,7). However, the extent to which these pathways are shared or differentially engaged across these conditions remains unclear.

Extracellular vesicles (EVs) - membrane-bound particles released by most cell types - play key roles in intercellular communication by transporting proteins, lipids, nucleic acids and other molecules (8,9). Beyond their function as mediators of intercellular communication, EVs and their contents reflect the physiological and pathological state of their cells of origin. As such, EVs provide a unique opportunity to capture integrated biological signals across multiple cellular compartments, offering a systems-level perspective on disease-associated processes.

Despite growing interest in the implications of EVs in chronic and inflammatory conditions, most studies have focused on an individual disease or the presence of isolated markers (10–12). Comparative analyses across clinically overlapping conditions remain limited, and the extent to which EV profiles can reveal coordinated alterations across interconnected biological pathways remains poorly defined. In fact, the evaluation of platelet-, immune-, or vascular-associated EV populations is just starting to be explored across the disorders examined in this study (11,13).

In this study, we adopted an integrative approach to small EV profiling by combining nanoparticle tracking analysis (NTA) with targeted detection of EV-associated surface markers co-expressed with CD9 by the implementation of AlphaScreen technology (28,30). We focused on small EVs because they constitute the predominant circulating EV population, are enriched in biologically active cargo, and have emerged as promising liquid biopsy biomarkers due to their stability and ability to reflect the physiological state of their cells of origin. Limiting analyses to this relatively homogeneous EV subset improves analytical reproducibility and facilitates the identification of disease-associated molecular signatures. The selected marker panel was designed to capture distinct yet interconnected biological processes involving EV-mediated intercellular communication. Platelet activation was assessed via CD41/CD61 (integrins αIIb and β3, forming the GPIIb/IIIa complex), the canonical marker pair for platelet-derived EVs, released upon platelet activation and implicated in thrombo-inflammatory crosstalk (14,15). Immune signaling was captured through CD74, the MHC class II invariant chain and functional receptor for macrophage migration inhibitory factor (MIF), which links antigen presentation machinery to MIF-driven pro-inflammatory and pro-survival signaling across immune and stromal cell types (16). Leukocyte trafficking was assessed via CX3CR1, the fractalkine receptor that mediates adhesion and migration of non-classical monocytes, NK cells, and effector/memory CD8+ T cells along the vascular endothelium. In plasma, CX3CR1-positive EVs are best interpreted as reflecting activity of these circulating, adhesion-competent, leukocyte subsets. In addition, CX3CR1 constitutes a marker of microglial activation. Angiogenesis and tissue remodeling were captured through CD13 (aminopeptidase N), an enzyme selectively upregulated on angiogenic (but not quiescent) vascular endothelium, which contributes to extracellular matrix degradation and cell migration during neovascularization, while also serving as a myeloid lineage marker (17). Vascular regulation through the renin-angiotensin system was assessed using ACE1 (CD143) and ACE2, which represent the two counter-regulatory arms of the RAS: ACE1 generates the vasoconstrictive, pro-inflammatory angiotensin II, while ACE2 converts angiotensin II to the vasoprotective angiotensin-(1–7), such that their combined profiling reflects the balance between these opposing vascular tone pathways (18,19). Finally, CD63, a tetraspanin enriched in intraluminal vesicles of multivesicular bodies, was included as a general EV-associated marker consistent with MISEV guidelines, serving as a lineage-independent confirmation of EV identity rather than a marker of a specific biological process (20).

By directly comparing EV-enriched plasma samples from ME/CFS, FM, COD, and LC-19 patients, and healthy controls (HCs), we aimed to identify disease-associated small EV signatures and evaluate their potential to distinguish clinically overlapping conditions. Furthermore, we sought to determine whether EV-associated markers could reveal dysregulation across platelet, immune, and vascular pathways, providing insight into the involved mechanisms and systemic nature of these disorders.

## METHODS

### Study cohort and ethical approval

This cross-sectional observational study was approved by the Public Health Research Ethics Committee DGSP-CSISP of Valencia (number: 20230602/11/01), Valencia, Spain. All participants provided written informed consent prior to inclusion in the study and were instructed to avoid medication intake for at least 12 hours prior to blood collection when applicable.

Platelet-free plasma samples were obtained from 50 female participants (n = 10 per group): ME/CFS, FM, COD, LC-19, and age-matched HCs. Samples and data from patients included in this study were provided by the Biobank IBSP-CV, integrated in the ISCIII Biobanks and Biomodels Platform and in the Valencian Biobanking Network and they were processed following standard operating procedures with the appropriate approval of the Ethics and Scientific Committees.

Patients were diagnosed according to established clinical criteria: Canadian Consensus Criteria (CCC) and/or International Consensus Criteria (ICC) for ME/CFS, American College of Rheumatology (ACR) 2010/2011 criteria for FM(1,2,4,21–23), and LC-19 patients were diagnosed according to current clinical guidelines (3,6). Clinical status and symptom severity were further assessed using standardized questionnaires, including the Multidimensional Fatigue Inventory (MFI) (24), the Short Form-36 (SF-36) (25), and the Fibromyalgia Impact Questionnaire (FIQ) (26). HC samples were selected from the IBSP-CV biobank collection to match patient age and sex; and were screened to exclude individuals with a history of chronic pain, fatigue-related disorders, or other relevant medical conditions. Individuals under medication or presenting any significant health condition were excluded.

All blood samples were collected and processed under standardized conditions by the Valencian IBSP-CV Biobank. To minimize ex vivo platelet activation and spurious release of platelet-derived vesicles, whole blood was processed fresh, with no freeze-thaw cycles prior to plasma separation, using a double-centrifugation protocol consisting of an initial low-speed spin (1,500 × g, 10 minutes, RT) to remove cells, followed by a second high-speed spin of the supernatant (2,500 × g, 10 minutes, RT) to deplete residual platelets, in line with the recommendations of the Scientific and Standardization Committee (SSC) of the International Society on Thrombosis and Hemostasis (ISTH) for pre-analytical standardization of plasma microparticle/EV studies. Platelet-poor plasma was isolated, aliquoted, and stored at −80°C until further analysis. Protocol adapted from ref Lacroix R et al., (27).

### Extracellular vesicle isolation

EVs were isolated from human plasma using Total Exosome Isolation Reagent from the plasma kit (Invitrogen, cat. n. 4484450) according to the manufacturer’s instructions, without proteinase treatment. Plasma samples were thawed and clarified by centrifugation at 2000 × g for 20 min at room temperature and then passed through a 0.22 µm polyethersulfone filter (Thermo Fisher Scientific) to remove residual cellular debris and large particles. While this step was included to improve sample homogeneity, this filtering may also partially exclude larger vesicles. Filtered plasma was transferred to a clean tube, diluted with 0.5 volumes of phosphate-buffered saline (PBS), and vortexed to mix thoroughly. Subsequently, 0.2 volumes of Exosome Precipitation Reagent (relative to the total plasma + PBS volume) were added, and the mixture was homogenized by gentle inversion until a uniform, slightly turbid solution was obtained. Samples were incubated at room temperature for 10 min to allow EV precipitation, then centrifuged at 10,000 × g for 5 min at room temperature. The supernatant was carefully removed by pipetting, leaving the EV-containing pellet at the bottom of the tube. An additional brief centrifugation step (10,000 × g, 30 s) was performed to collect residual reagent prior to complete supernatant removal. EV pellets were resuspended in PBS (25 µL for 100 µL starting plasma) by gentle pipetting. Resuspended EVs were stored at −80°C for downstream analyses. To standardize sample processing and ensure consistency across measurements, EV suspensions underwent only one freeze-thaw cycle prior to their analyses. EVs were isolated by polymer-based precipitation, a method that co-precipitates lipoproteins and protein aggregates that cannot be resolved from vesicles. In accordance with MISEV2023, the resulting preparations are therefore referred to throughout as EV-enriched fractions, as co-isolation of non-vesicular components cannot be excluded.

### Nanoparticle tracking analysis (NTA)

EV size distribution and particle concentration were measured by NTA using a NanoSight NS300 instrument (Malvern Panalytical, Malvern, UK) equipped with NTA software version 3.2 Dev Build 3.2.16. Samples were diluted in filtered PBS to achieve optimal particle concentration within the recommended detection range. For each sample, five independent recordings of 30 seconds were acquired using a standardized protocol. Acquisition settings were as follows: sCMOS (scientific complementary metal–oxide–semiconductor) camera, green laser, camera level 11, shutter 600, gain 300, frame rate 25 frames per second, and 749 frames per capture. Measurements were performed at 25°C with a viscosity setting of 0.9 centipoise cP. Analysis parameters included a detection threshold of 10, automatic blur size, and automatic maximum jump distance (14.4-15.2 pixels). Particle size and concentration were calculated based on Brownian motion using the Stokes-Einstein equation (28). Particles within the 40-200 nm size range were considered representative of small EVs and were used for downstream normalization (8).

### EV-associated protein detection by AlphaScreen

EV-associated surface proteins were analyzed using AlphaScreen-based proximity immunoassays (PerkinElmer/BMG Labtech technology), based on the amplified luminescent proximity homogeneous assay principle (AlphaScreen), also known as ExoScreen (29,30).

In this assay EV-associated antigens are detected through dual antibody recognition. A biotinylated antibody binds to the streptavidin-coated donor beads (PrekinElmer, cat. n. 6760002), while a second anti-CD9 antibody (Cosmo Bio Co., LTD, cat. n. SHI-EX0-M01-50) recognizing a different epitope is conjugated to the unconjugated AlphaLISA acceptor beads (PrekinElmer, cat. n. 6772001). Upon excitation at 680 nm, donor beads generate singlet oxygen, which induces light emission at 615 nm in nearby acceptor beads when both are within ∼200 nm, enabling sensitive detection of vesicle-associated proteins (31).

EV-enriched samples were incubated with biotinylated antibodies and acceptor bead-conjugated antibodies in a 384-well AlphaPlate (PerkinElmer, cat. n. 6005350). Plates were incubated overnight at 4°C to allow binding of EV-associated antigens to the antibody complexes. The following day, plates were equilibrated to room temperature under dark conditions, and streptavidin-coated donor beads were added to each well (final reaction volume: 27.5 µL per well). After gentle spin to collect contents at the bottom of the wells, plates were incubated for 30 minutes at room temperature in the dark to allow proximity-dependent signal generation. Luminescence signal counts were measured using a CLARIOstar multimode plate reader (BMG LABTECH) under AlphaScreen settings. All samples were measured in technical duplicates, and care was taken to minimize light exposure throughout the assay.

A panel of EV surface markers was selected to characterize distinct vesicle subpopulations. CD63 (Miltenyi Biotec, cat. n. 130-136-549) was used as general EV-associated marker and for minimal characterization of EV-enriched fractions in accordance with MISEV2023 recommendations. Additional markers were selected to capture complementary biological processes. Immune-associated EVs were assessed using CD74 (Milteny Biotec, cat. n. 130-119-033), related to antigen presentation and macrophage migration inhibitory factor (MIF) signaling, and CX3CR1 (Milteny Biotec, cat. n. 130-132-764), a chemokine receptor involved in leukocyte adhesion and trafficking. Renin-angiotensin system-associated EVs were evaluated through ACE1 (or CD143) (Milteny Biotec, cat. n. 130-107-950), associated with pro-inflammatory and vasoconstrictive signaling, and ACE2 (Stemcell Technologies, cat. n. 100-0761), linked to counter-regulatory, vasoprotective pathways. Myeloid- and platelet-associated EVs were characterized using CD13 (Miltenyi Biotec, cat. n. 130-119-572), a membrane aminopeptidase involved in tissue remodeling and angiogenesis, and CD41/CD61 (Miltenyi Biotec, cat. n. 130-109-421), components of the platelet GPIIb/IIIa complex indicative of platelet activation.

All incubations and handling steps were performed under low-light conditions to prevent signal degradation.

### Data processing and statistical analysis

Data processing and statistical analyses were performed using a combination of Python (version 3.12.2) and R (version 2024.09.0+375). Python was primarily used for data preprocessing, whereas statistical testing, receiver operating characteristic (ROC) analysis and data visualization were mainly conducted in R using packages including *ggplot2*, *dplyr* and *pROC*.

NTA-derived particle concentration and size distribution data were processed to obtain mean particle size and total particle counts. For downstream analyses, particles within the 40-200 nm size range were considered representative of the small EV population (8). AlphaScreen luminescence signals were normalized to EV particle concentration within the 40-200 nm size range to account for inter-sample variability.

Comparisons between experimental groups (ME/CFS, FM, COD, LC-19, and HC) were performed using the non-parametric Wilcoxon rank-sum test (also known as Mann-Whitney U test). Data visualization was primarily performed in R using box and violin plots with overlaid individual data points to represent group distributions. Outlier values were identified within each group using Tukey’s rule (1.5 × interquartile range) and excluded from the analysis.

ROC curve analysis was conducted to evaluate the discriminatory capacity of EV marker profiles between LC-19 group and remaining groups (ME/CFS, FM, COD and HC). The area under the curve (AUC) was calculated as a measure of classification performance.

Correlation analyses between marker signals and clinical variables were performed in Python using SciPy. Because the AlphaScreen plate layout provided two technical replicate measurements per participant, replicate wells were first averaged so that each patient contributed a single value per marker, and all correlations were computed at the patient level (n = 10 per group). Associations were assessed using Spearman’s rank correlation coefficient (R), which does not assume linearity or normality. For each marker-variable pair, coefficients were calculated using pairwise complete observations and exact two-sided p-values were obtained from the corresponding rank correlation test. Analyses were run separately within each disease group and covered the full set of variables available for every participant: 37 routine blood parameters and 17 FIQ, MFI and SF-36 subdomains. Higher scores on the FIQ and MFI indicate greater symptom burden, whereas higher scores on the SF-36 indicate better health status or function; correlation directions are reported with respect to these scoring conventions. Correlation coefficients, p-values, and the number of observations used for each pairwise comparison were compiled into matrices and visualized as heatmaps, with colour gradients representing the strength and direction of the correlations and asterisks indicating associations that reached significance: * (p ≤ 0.05), ** (p ≤ 0.01), *** (p ≤ 0.001).

## RESULTS

### Study cohort

A total of 50 female participants (n= 10/study group) were included in the study, comprising five groups: ME/CFS, FM, COD, LC-19, and HC. Patients with ME/CFS and FM were clinically diagnosed according to established clinical criteria (CCC and/or ICC) for ME/CFS, 2010/2011 ACR criteria for FM (**Table 1**), while LC-19 patients met current clinical definitions of post-COVID condition based on persistent symptoms following severe acute respiratory syndrome coronavirus 2 (SARS-CoV-2) infection (3,6). The COD group included individuals fulfilling diagnostic criteria for both ME/CFS and FM (**Table 1**). HC were selected to match patient groups and had no history of chronic pain, fatigue-related disorders, or other relevant medical conditions.

**Table 1.** Patient demographics and health status assessment with FIQ, MFI, and SF-36 (24–26) questionnaires (n = 50; 10/study group). Data are presented as the mean +-standard deviation (SD), and range values.

|  | **ME/CFS (n=10)** |  |  | **FM (n=10)** |  |  | **COD (n=10)** |  |  | **LC-19 (n=10)** |  |  |
|---|---|---|---|---|---|---|---|---|---|---|---|---|
|  | **Mean** | **SD** | **Range** | **Mean** | **SD** | **Range** | **Mean** | **SD** | **Range** | **Mean** | **SD** | **Range** |
| **Demographics** |  |  |  |  |  |  |  |  |  |  |  |  |
| Age | 54.6 | 15.99 | (19 – 68) | 55.22 | 11.22 | (27 – 62) | 52.13 | 5.46 | (47 – 64) | 47.1 | 5.80 | (38 – 54) |
| BMI | 29.47 | 5.05 | (21 - 39.5) | 27.54 | 4.62 | (22.4 - 36.4) | 26.99 | 6.20 | (18 - 37.5) | N/A | N/A | N/A |
| Years since disease diagnosis* | 13 | 8.60 | (1 – 28) | 11.6 | 13.79 | (1 – 45) | 10.2 | 8.87 | (1 – 26) | 2 | 1.49 | (0 – 5) |
| Age when diagnosed | 41.6 | 14.77 | (17 – 62) | 42.89 | 15.00 | (17 – 57) | 40.38 | 8.63 | (25 – 54) | 45.1 | 6.64 | (35 – 53) |
| **FIQ Questionnaire** |  |  |  |  |  |  |  |  |  |  |  |  |
| Total FIQ | 69.99 | 14.93 | (41.32-90.62) | 69.55 | 21.09 | (18.44-89.88) | 77.76 | 9.62 | (63.23-89.63) | 61.12 | 20.24 | (21.43-92.51) |
| Function | 5.64 | 2.24 | (1.65-8.25) | 5.15 | 2.74 | (0-9.9) | 7.33 | 1.78 | (3.3-9.9) | 5.02 | 2.89 | (0-9.57) |
| Overall | 8.44 | 1.25 | (7.15-10.01) | 7.87 | 1.94 | (4.29-10.01) | 6.51 | 1.26 | (4.29-8.58) | 6.51 | 2.78 | (0-8.58) |
| Symptoms | 8.01 | 2.71 | (1.43-10.01) | 5.43 | 2.5 | (1.43-8.58) | 7.47 | 2.12 | (4.29-10.01) | 6.36 | 3.36 | (1.43-10.01) |
| **MFI Questionnaire** |  |  |  |  |  |  |  |  |  |  |  |  |
| General Fatigue | 17.7 | 3.74 | (8-20) | 17.4 | 3.5 | (11-20) | 19.11 | 2.03 | (14-20) | 18 | 2.36 | (14-20) |
| Physical Fatigue | 16.4 | 4.65 | (4-20) | 16.8 | 3.26 | (11-20) | 18.56 | 2.3 | (14-20) | 18.1 | 2.13 | (14-20) |
| Reduced Activity | 14 | 4.78 | (4-20) | 17.5 | 3.27 | (10-20) | 17.78 | 3.38 | (10-20) | 16.9 | 2.77 | (13-20) |
| Reduced Motivation | 12.1 | 4.33 | (5-20) | 11 | 4.29 | (4-19) | 14.78 | 3.7 | (10-20) | 13 | 4.16 | (8-19) |
| Mental Fatigue | 13.8 | 5.35 | (4-20) | 16.6 | 3.44 | (9-20) | 18.11 | 2.71 | (13-20) | 15.1 | 4.93 | (6-20) |
| **SF-36 Questionnaire** |  |  |  |  |  |  |  |  |  |  |  |  |
| Physical Functioning | 30.5 | 13.83 | (10-50) | 32.5 | 16.03 | (5-45) | 26.11 | 16.54 | (0-50) | 50.56 | 26.86 | (5-85) |
| Role Physical | 8.2 | 8.1 | (0.75-25.75) | 11.6 | 5.96 | (7.25-21) | 3.89 | 5.07 | (0.75-14.75) | 8.31 | 7.29 | (0.75-21.75) |
| Bodily Pain | 39 | 21.61 | (10-77.5) | 35 | 22.58 | (10-80) | 27.22 | 16.7 | (10-55) | 50.83 | 26.98 | (10-87.5) |
| General Health | 30 | 12.25 | (10-50) | 24 | 12.43 | (5-35) | 11.67 | 11.18 | (0-30) | 27.78 | 15.23 | (5-45) |
| Vitality | 25.5 | 14.03 | (10-50) | 23.5 | 16.67 | (10-30) | 13.33 | 5.59 | (10-25) | 23.89 | 16.35 | (10-60) |
| Social Functioning | 33.5 | 28.9 | (0-90) | 37.5 | 32.55 | (0-90) | 38.06 | 21.64 | (0-70) | 54.72 | 25.99 | (10-90) |
| Role Emotional | 24.67 | 8.48 | (9.67-36.67) | 22.3 | 14.72 | (0.67-36.67) | 21.44 | 15.52 | (0.67-36.67) | 25.7 | 11.66 | (0.67-36.67) |
| Mental Health | 52.4 | 20.95 | (28-84) | 43.6 | 16.38 | (24-68) | 43.56 | 23.62 | (16-72) | 55.11 | 20.57 | (12-80) |

*Abbreviations: COD: co-diagnosis of ME/CFS and FM, FIQ – Fibromyalgia Impact Questionnaire, FM – Fibromyalgia, LC-19 – Long COVID-19, ME/CFS – Myalgic encephalomyelitis/chronic fatigue syndrome, MFI – Multidimensional Fatigue Inventory, SF-36 - Short Form-36, SD - standard deviation, N/A – Not Available.*

Patient demographics and instrument-based phenotyping are summarized in **Table 1** and detailed in **Supplementary Table S1**. As an overall, no significant differences were observed between groups (p > 0.05, FDR-corrected), indicating adequate matching across these variables, except for age differences between LC-19 and ME/CFS (Kruskal-Wallis, p = 0.046, FDR-corrected), with the LC-19 cohort being younger than the ME/CFS group (Dunn’s post hoc, p = 0.032). As expected, given the recent emergence of Long COVID, years since diagnosis also differed significantly across groups (Kruskal-Wallis, p = 0.032, FDR-corrected), with the LC-19 cohort showing a substantially shorter disease duration than the ME/CFS, FM, and COD groups (Dunn’s post hoc, all p ≤ 0.05).

No significant differences were observed between groups in any FIQ, MFI, or SF-36 subdomain scores (p>0.05), indicating a similar degree of severity across the disease groups studied here, except for SF-36 Physical Functioning and General Health p-values (p = 0.045, p = 0.020) which despite reaching significance, did not survive correction by either Kruskal Wallis or one way ANOVA FDR assessment. These findings indicate a broadly comparable symptom burden across the ME/CFS, FM, COD, and LC-19 groups. According to total FIQ all patient groups were considered severely affected (≥59) (**Table 1**).

### EV concentration and size distribution differ across disease group

To characterize the overall EV profile across study groups, particle concentration and size distribution (diameter in nm) were assessed by NTA, with five technical replicate measurements performed per sample (n = 10 samples/study group; 50 data points per group). EV-enriched fractions were detected in all samples, with particle size distributions predominantly within the expected range for small EV (40-200 nm in diameter). Within this size range distribution, the profiles revealed variability across groups (**Figure 1A**). Raw measures are available in **Supplementary Table S2**.

**Figure 1.**
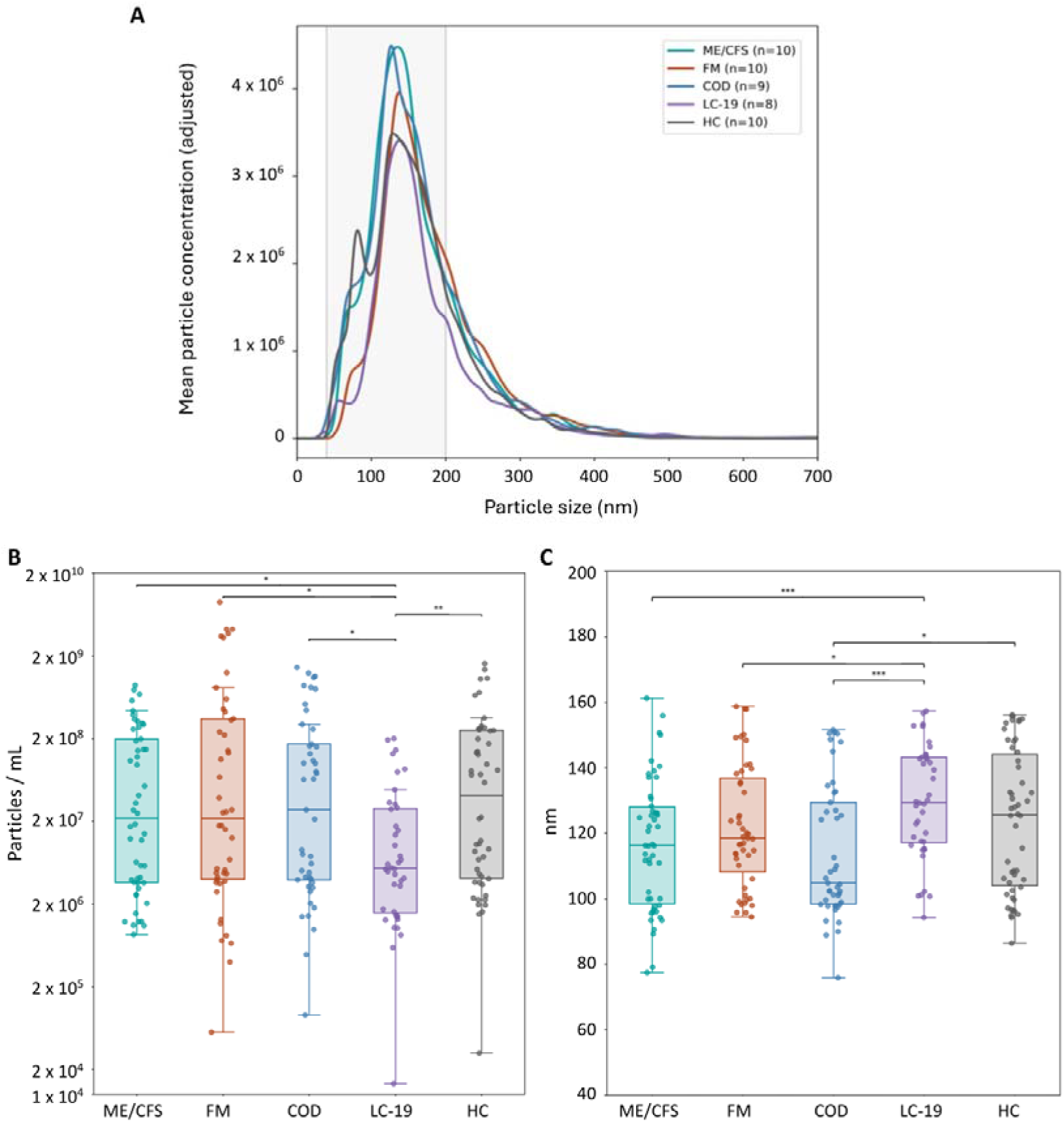
Particle size distribution and comparison of EV concentration and size across study groups. EVs subjected to a single freeze-thaw cycle were isolated as described in the Methods section. Five technical replicate measurements were performed per sample (n = 10 samples/study group; 50 data points per group). **A)** NTA particle size distribution profiles of EVs. Histogram overlay of the mean particle size distribution curves for all groups analyzed. The grey shaded area indicates diameter size range used for downstream analyses, corresponding to small EVs (40-200 nm). Particles outside this range were excluded to minimize potential contributions from larger vesicles, aggregates, or smaller particles that may not correspond to EV populations. **B)** Particle concentration (particles/mL of plasma) measured by NTA in the five study groups: ME/CFS, FM, COD, LC-19, and HC. **C)** Mean particle size (diameter in nm) measured by NTA in the same experimental groups. Statistical comparisons were performed using the Wilcoxon test indicated in the plot as * (p ≤ 0.05), ** (p ≤ 0.01), *** (p ≤ 0.001). Plots were generated using Python. *Abbreviations: COD: co-diagnosis of ME/CFS and FM, FM – Fibromyalgia, HC – Healthy control, LC-19 – Long COVID-19, ME/CFS – Myalgic encephalomyelitis/chronic fatigue syndrome*.

Analysis of particle concentration revealed that the LC-19 group exhibited significantly lower particle levels than all other groups (Mann-Whitney U test: ME/CFS vs LC-19 (p = 0.012), FM vs LC-19 (p = 0.018), COD vs LC-19 (p = 0.013), and LC-19 vs HC (p = 0.002)), indicating reduced circulating EV levels associated with this condition (**Figure 1B)**. Analysis of mean particle size (diameter in nm) showed comparable size profiles for LC-19 patients and HC (p = 0.2) (**Figure 1C**); while, EVs from the other three disease groups (ME/CFS, FM, and COD) exhibited smaller mean particle than LC-19 (Mann-Whitney U test: ME/CFS vs LC-19 (p = 9.92×10), FM vs LC-19 (p = 0.030), and COD vs LC-19 (p = 6.25×10) COD EVs also differed significantly from HC (p = 0.037). Together, these findings suggest potential disease-specific EV profiles. All quantitative analysis was performed in five technical replicates. See the **Supplementary Table S3 and S4** for raw data.

### CD9/CD63-positive EVs exhibit similar levels across the analyzed groups

The overall abundance of EVs carrying canonical tetraspanin markers, CD9/CD63-positive vesicles were quantified by AlphaScreen assays, using two technical replicates per sample (n = 10 samples/study group; 20 data points/study group). The final number of observations after outlier exclusion (Tukeýs rule criteria) are indicated in the figure legend. CD9/CD63-positive EVs were consistently detected across all study groups (**Figure 2**), with no statistically significant differences observed between ME/CFS, FM, COD, LC-19, and HC. This indicates that, despite differences observed in overall particle concentration, the total pool of CD9/CD63-positive EVs remains relatively stable across conditions. However, an elevated tendency was noticed in LC-19 when compared to the HC group (fold change = 1.26; p = 0.1). For statistical analysis details see **Supplementary Table S5**.

**Figure 2.**
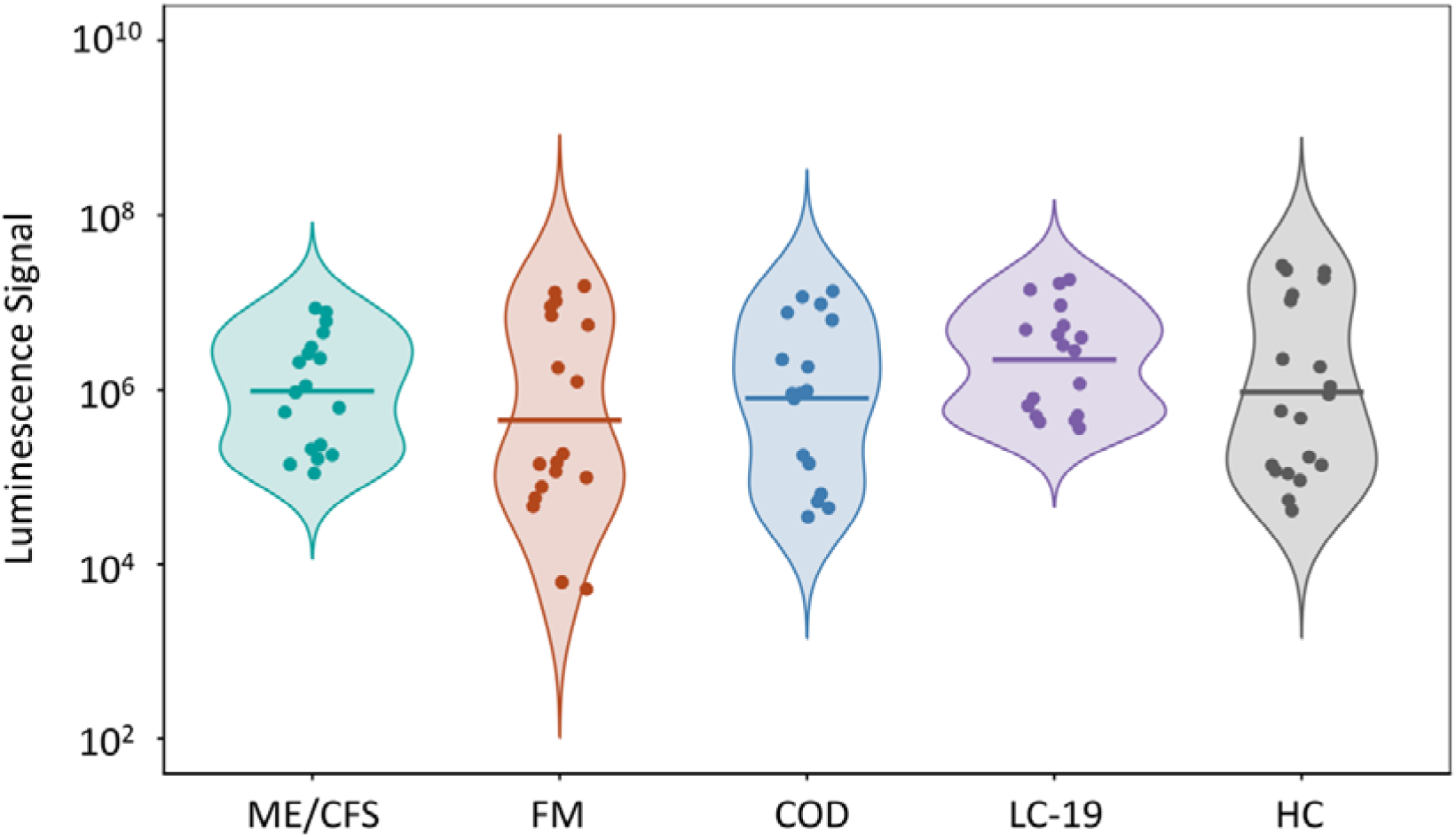
CD9/CD63-positive EV levels across study groups. Plasma EVs co-expressing CD9 and CD63 from ME/CFS (n=18), FM (n=18), COD (n=18), LC-19 (n=18) patients and HC (n=20) were assessed by AlphaScreen. Signal intensity reflects the abundance of EVs carrying both markers within each sample. AlphaScreen luminescence signals were normalized to the concentration of EV particles within the 40-200 nm size range measured by nanoparticle tracking analysis. Statistical significance determined by Wilcoxon test * (p ≤ 0.05), ** p ≤ 0.01, *** p ≤ 0.001. *Abbreviations: COD: co-diagnosis of ME/CFS and FM, FM – Fibromyalgia, HC – Healthy control, LC-19 – Long COVID-19, ME/CFS – Myalgic encephalomyelitis/chronic fatigue syndrome*.

### Immune-associated EV subpopulations are selectively enriched in LC-19

To investigate disease-associated EV phenotypes, immune-related EV subpopulations were analyzed based on co-expression of CD9 with CD74 and CX3CR1. In both assays, two technical replicates AlphaScreen were obtained per sample (n = 10 samples/study group; 20 data points/study group). The total number of observations after outliers’ removal (Tukeýs rule criteria) is shown in **Figure 3**. LC-19 samples exhibited significantly higher levels of CD9/CD74-positive EVs compared to all other groups (LC-19 compared to ME/CFS (p = 0.002), FM (p = 0.001), and COD (p = 0.029), as well as compared to HC (p = 0.021)) (**Figure 3A**), for more details see **Supplementary Table S6**. Similarly, CD9/CX3CR1-positive EVs were increased in LC-19, with statistically significant differences observed relative to HCs, FM, and ME/CFS groups [LC-19 compared to ME/CFS (p = 0.008), FM (p = 0.035), and HC (p = 0.002), and an elevated tendency for the COD group (p = 0.075)] (**Figure 3B**), see **Supplementary Table S7** for more detailed information. These results indicate a selective enrichment of immune-associated EV subpopulations in LC-19, consistent with altered immune signaling in this condition.

**Figure 3.**
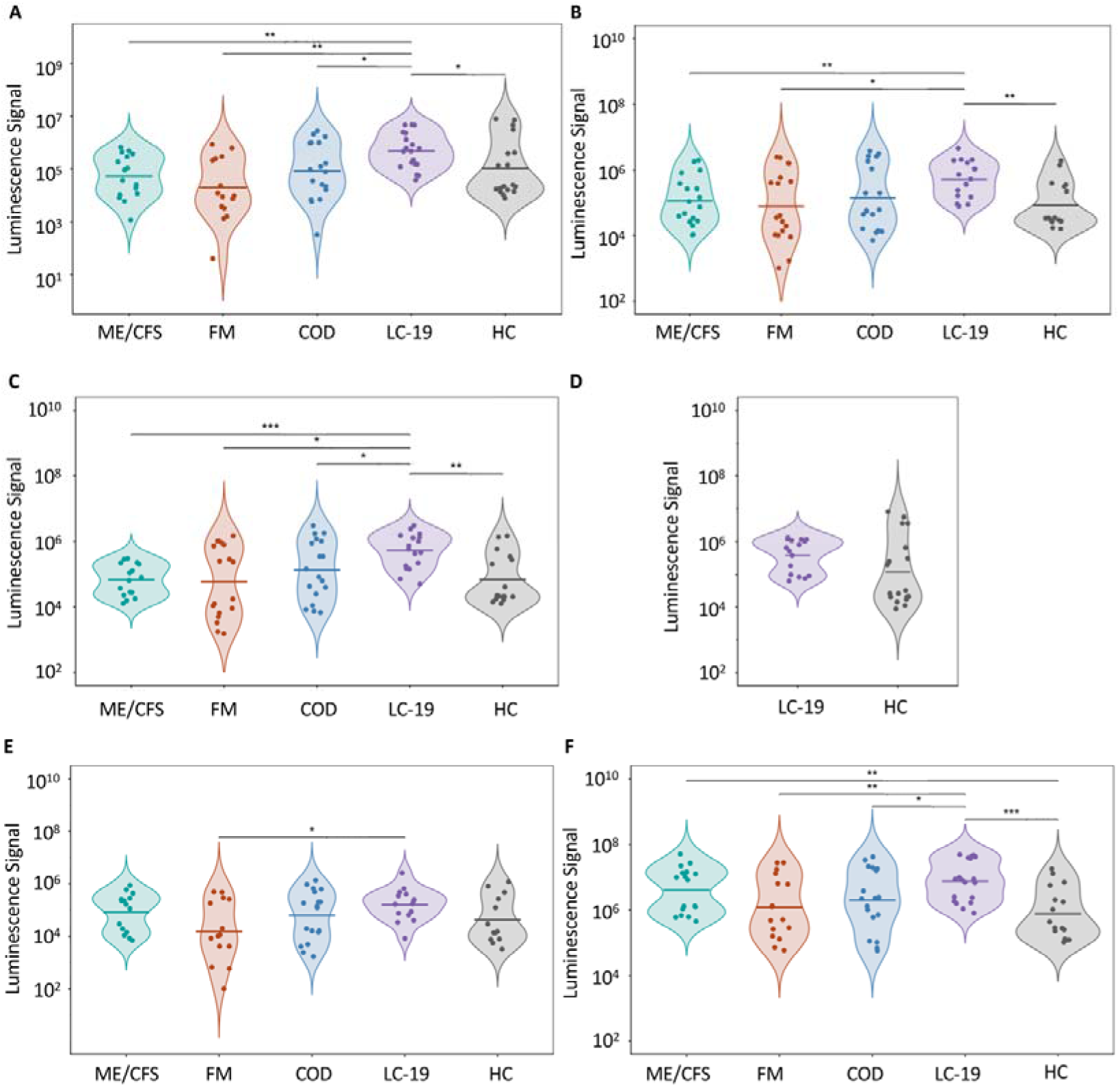
Immune-, renin-angiotensin system-, and myeloid/platelet-associated extracellular vesicle subpopulations in plasma. Plasma EVs from ME/CFS, FM, COD, and LC-19 patients and HC (n= 20 subjects/group; 100 observations) were assessed via AlphaScreen assays to detect the co-expression of **A)** CD9 and CD74 (ME/CFS, n = 17; FM, n = 16; COD, n =18; LC-19, n = 19; HC, n = 19), **B)** CD9 and CX3CR1 (ME/CFS, n = 18; FM, n = 18; COD, n =18; LC-19, n = 16; HC, n = 16), **C)** CD9 and ACE1(ME/CFS, n = 16; FM, n = 17; COD, n =18; LC-19, n = 17; HC, n = 17), **D)** CD9 and ACE2 (LC-19, n = 16; HC, n = 18), **E)** CD9 and CD13 (ME/CFS, n = 15; FM, n = 14; COD, n =18; LC-19, n = 15; HC, n = 13), and **F)** CD9 and CD41/CD61 (ME/CFS, n = 17; FM, n = 15; COD, n =18; LC-19, n = 19; HC, n = 15). Statistical analyses were performed using the Wilcoxon test indicated in the plot as * (p ≤ 0.05), ** (p ≤ 0.01), *** (p ≤ 0.001). *Abbreviations: COD: co-diagnosis of ME/CFS and FM, FM – Fibromyalgia, HC – Healthy control, LC-19 – Long COVID-19, ME/CFS – Myalgic encephalomyelitis/chronic fatigue syndrome*.

### RAS-associated EVs display selective enrichment in LC-19 plasma

Given the role of the RAS in vascular and inflammatory regulation, EVs expressing ACE1 (also known as CD143) and ACE2 were quantified. Two technical replicate measurements per sample were obtained (n = 20/study group). Similar to previous assessments, outliers were removed for downstream analysis. CD9/ACE1-positive EVs were significantly increased in LC-19 compared to all other groups (LC-19 vs ME/CFS (p = 2.1×10), LC-19 vs FM (p = 1.4 ×10 2), LC-19 vs COD (p = 4.8 ×10 ^2^) and LC-19 vs HC (p = 2.0 ×10 ^3^) (**Figure 3C**), showing a marked enrichment of ACE1-associated vesicles which suggests a shift of the renin-angiotensin system in these patients. To gain further insight into the relationship between ACE2 and ACE1 in LC-19, and considering the pivotal role of ACE2 as the entry receptor for SARS-CoV-2 (56–58), we measured circulating CD9/ACE2-positive EV levels in LC-19 patients and compared them with those observed in HC. Unexpectedly, the results showed slight but also higher levels of ACE2 in LC-19, which despite not reaching statistical significance (p = 0.07) showed a tendency (median fold change 20-fold higher in LC-19 vs. HC) (**Figure 3D**) (**Supplementary Table S8 and S9**). Although the literature describes ACE1 function related to vasoconstriction, and ACE2 associating to vasodilation (59,60), it is possible that compensatory mechanisms exist or that differences in cells are not captured by CD9-positive EVs.

### Myeloid/endothelial- and platelet-associated EVs display study group-specific alterations

Given the role of CD13 (aminopeptidase N) in angiogenesis and myeloid lineage identity and the role of the CD41/CD61 (GPIIb/IIIa) complex in platelet activation, myeloid- and platelet-associated EV subpopulations were analyzed based on the co-expression of CD9 with CD13 and CD41/CD61. Once more, two technical replicates were obtained for all study group EVs (n = 50; 100 AlphaScreen assays). As in all other analysis, identified outliers were excluded by applying Tukey’s rule. A significant increase in the number of CD9/CD13-positive EVs was only observed in the LC-19 group compared with the FM group (p = 0.04) (**Figure 3E and Supplementary Table S10**), pointing at this characteristic as a potential differentiator of these two patient groups. Meanwhile, a significant increase in CD9/CD41/CD61-positive EVs was detected in the ME/CFS and LC-19 groups compared with the HC group, and in the LC-19 group compared with the FM and COD groups (ME/CFS vs HC: p = 0.008, FM vs COD: p = 0.47, FM vs LC-19: p = 0.007, FM vs HC: p = 0.45, COD vs LC-19: p = 0.048, COD vs HC: p = 0.2, and LC-19 vs HC: p = 0.0009) (**Figure 3F**). Raw data available as the **Supplementary Table S11**.

These results point to differential involvement of myeloid- and platelet-associated EVs across disease groups, with a particularly pronounced involvement of both ME/CFS and LC-19 conditions.

### EV marker profiles discriminate LC-19 from other groups

To evaluate the potential of EV-associated markers as disease-discriminative features, ROC analysis was performed (n = 50; 10/study group).

Individual EV surface markers showed variable discriminative capacity for LC-19 versus the other groups (**Figure 4A**): CD63 (AUC = 0.642), CD74 (AUC = 0.764), CX3CR1 (AUC = 0.739), ACE1 (AUC = 0.777), ACE2 (AUC = 0.684), CD13 (AUC = 0.650), and CD41/61 (AUC = 0.721). Combined EV marker signatures distinguished LC-19 patients from healthy controls and other disease groups with high accuracy (Figure 4B), as reflected by an AUC of 0.914. These results support the potential utility of EV-associated signatures for distinguishing LC-19 from related conditions. See **Supplementary Table S12**.

**Figure 4.**
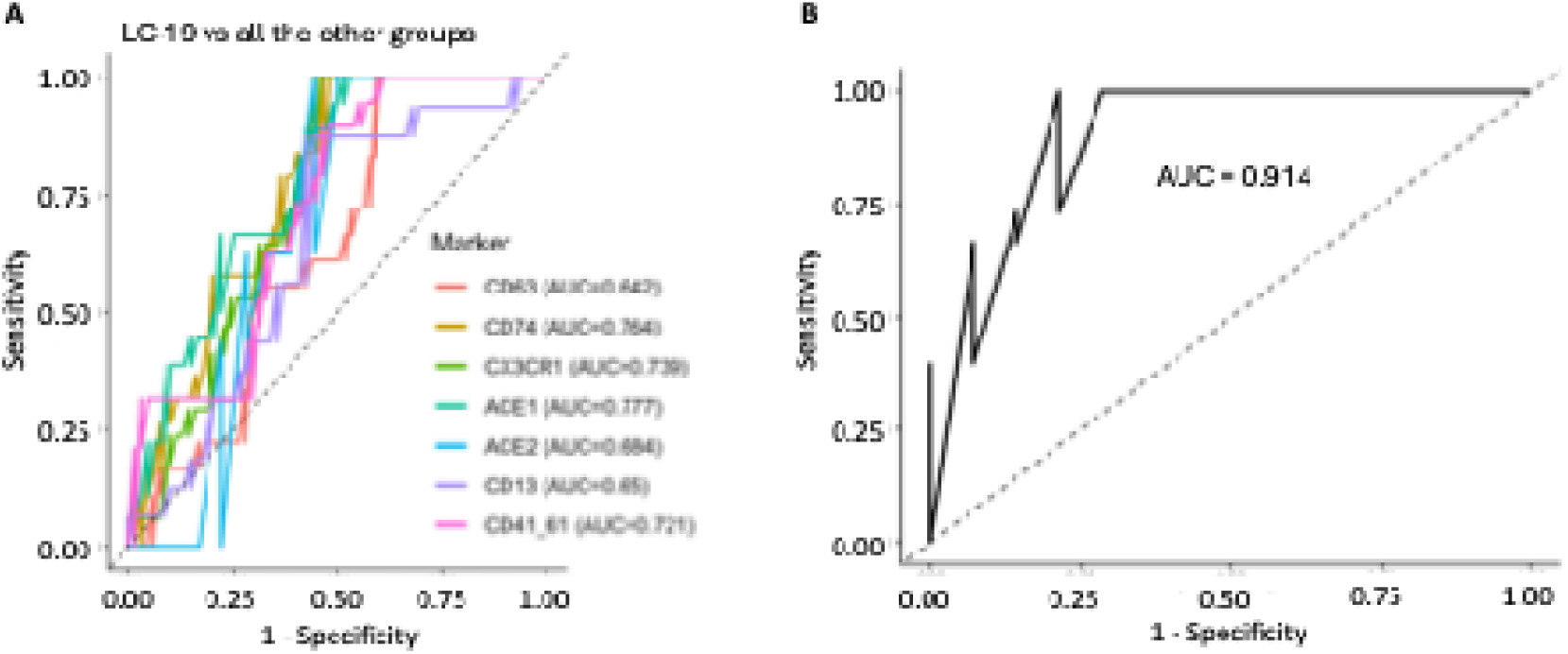
ROC-based evaluation of EV-associated markers for LC-19 classification. ROC curves to evaluate the discriminative potential of EV-associated markers between LC-19 patients and all the other groups (HC, ME/CFS, FM, and COD) (n= 10 subjects/group) are shown. **A)** ROC curves for individual EV markers, with AUC values ranging from 0.6 to 0.8, indicating individual marker discriminative performance are displayed. **B)** ROC analysis of the combined EV marker panel (all markers). *Abbreviations: COD: co-diagnosis of ME/CFS and FM, FM – Fibromyalgia, HC – Healthy control, LC-19 – Long COVID-19, ME/CFS – Myalgic encephalomyelitis/chronic fatigue syndrome*.

### EV marker profile correlates with symptomatology and blood parameters

Spearman rank correlation analysis was used to assess monotonic associations between marker luminescence intensities and clinical variable values (instrument scores and baseline clinical blood analytics). Replicate AlphaScreen measurements were averaged so that each participant contributed a single value per marker, and only variables available for all participants were included in the analysis. Correlation coefficients, p-values and the number of observations available for each pairwise comparison is displayed visualized as heatmaps (**Figures 5 and 6**, Supplementary Table S13).

**Figure 5.**
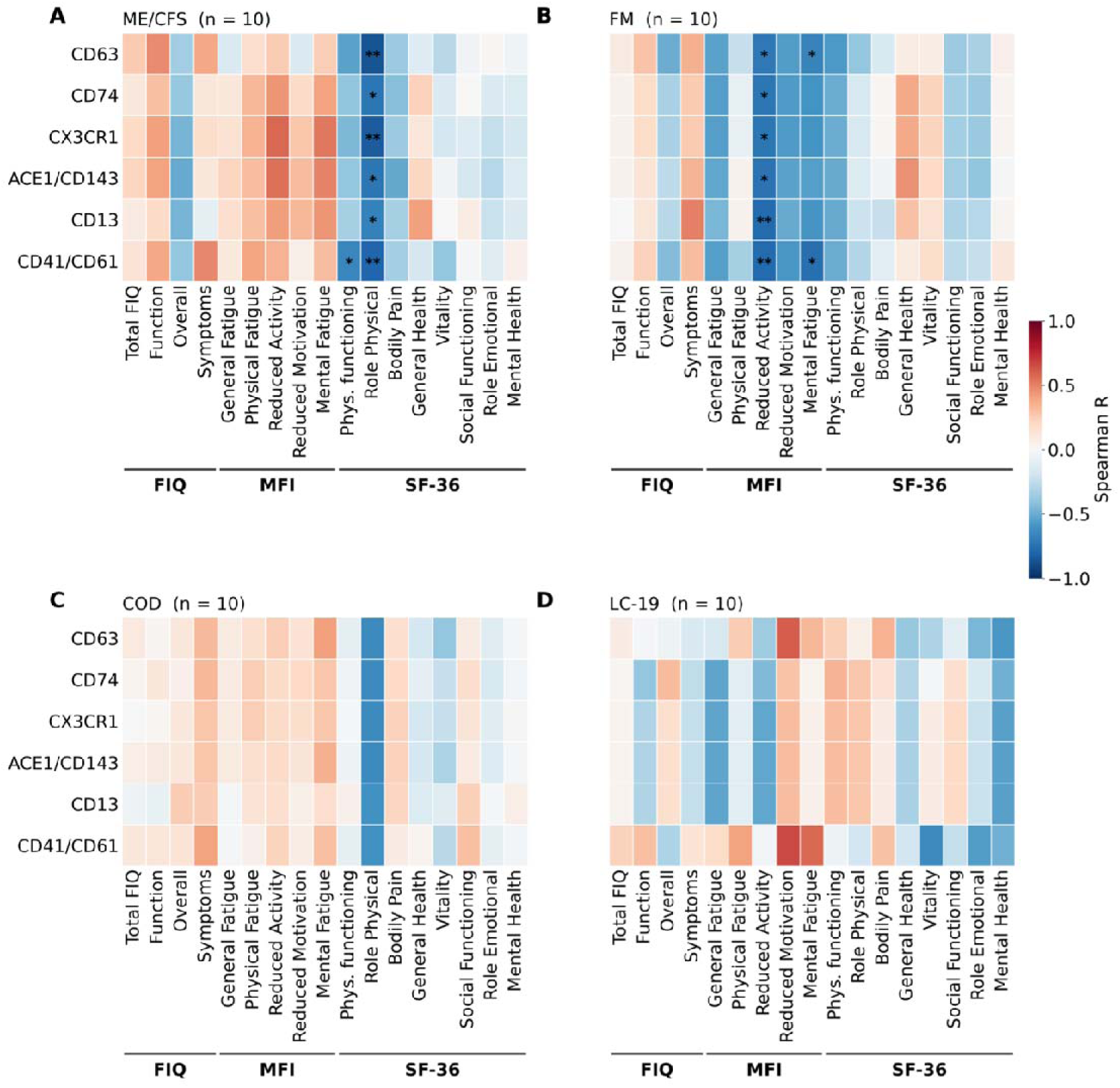
Correlation heatmaps of EV markers and patient-reported symptoms. Heatmaps display rank correlation coefficients between the EV surface marker profiles and the FIQ, MFI and SF-36 domain scores, computed separately within each patient group at the patient level (n = 10 per group). **A)** ME/CFS, **B)** FM, **C)** COD, **D)** LC-19. Domains are grouped beneath each panel by questionnaire (FIQ, MFI and SF-36). Colour indicates the direction and strength of the association and asterisks mark the pairs reaching significance: * (p ≤ 0.05), ** (p ≤ 0.01).

**Figure 6.**
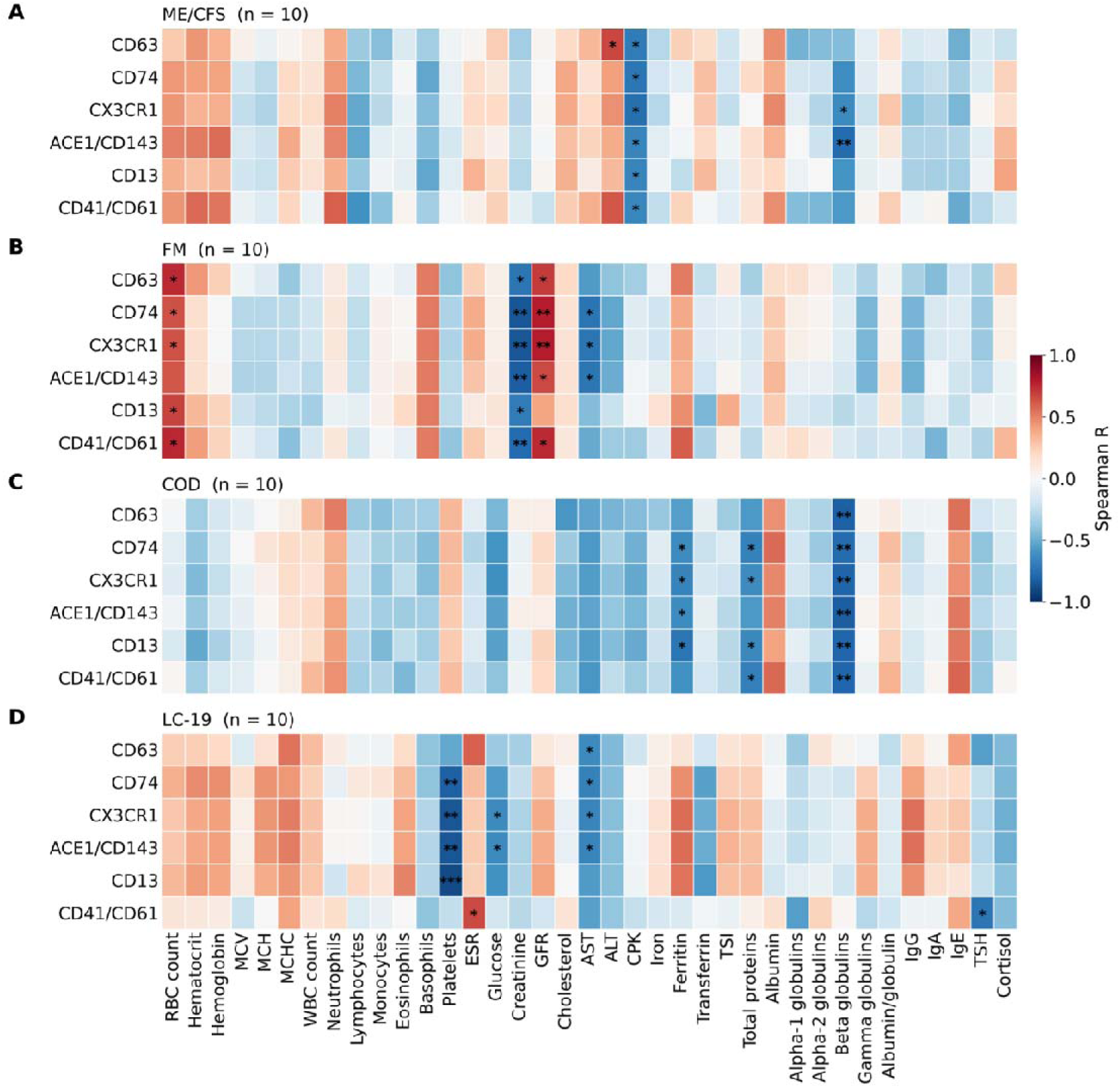
Correlation heatmaps of EV markers and patient-reported symptoms and clinical routine blood parameters. Heatmaps show Spearman correlation coefficients between the EV surface markers and the measured baseline blood parameters. **A)** ME/CFS, **B)** FM, **C)** COD, **D)** LC-19. Colour indicates the direction and strength of the association and asterisks mark the pairs reaching significance: * (p ≤ 0.05), ** (p ≤ 0.01), *** (p ≤ 0.001).

Spearman correlation coefficients (R) between EV marker expression (CD63, CD74, CX3CR1, ACE1, CD13, CD41/CD61) and: (a) patient-reported symptom severity (FIQ, MFI, and SF-36); or (b) clinical blood parameters, are shown by color code and intensity (pink/red for positive correlations and blue for negative) for each patient group (ME/CFS, FM, COD, LC-19; n = 10 per group). It should be pointed that FIQ and MFI higher scores indicate greater symptom burden; while higher scores of SF-36 subdomains indicate improved health status/function.

Overall, the analysis identified heterogeneous, group-specific relationships between EV marker signals and clinical variables. The observation that only a few clinical variables were associated with multiple EV markers suggests that different EV populations may provide complementary insights into the pathophysiological processes into the mechanisms driving shared disease manifestations.

#### EV marker-symptom correlations

Main significant symptom correlations in the ME/CFS group (**Figure 5A**), were negatively associated with the SF-36 Role Physical subdomain [CD63 (R = −0.860, p = 0.0014); CX3CR1 (R = −0.829, p = 0.0030); CD41/CD61 (R = −0.793, p = 0.0062); CD74 and ACE1(CD143) (both R = −0.726, p = 0.0175); and CD13 (R = −0.677, p = 0.0316)], indicating that higher EV marker levels associate with greater physical health limitations. In addition, the EV marker CD41/CD61 also negatively correlated with SF-36 Physical Functioning subdomain (R = −0.654, p = 0.0400) indicating a predominant association of the platelet activation pathway with a worse health status in this patient subgroup.

While in the FM patient group correlations concentrated on two related fatigue domains (Figure 5B) for all six markers. This time the negative correlations affected the MFI Reduced Activity subdomain [CD13 and CD41/CD61 (both R = −0.774, p = 0.0086); ACE1(CD143) (R = −0.736, p = 0.0152); CD74 and CX3CR1 (both R = −0.724, p = 0.0180); and CD63 (R = −0.717, p = 0.0195)]. In addition, CD41/CD61 and CD63 levels negatively correlated with the MFI Mental Fatigue subdomain (R = −0.751, p = 0.0123 and R = −0.665, p = 0.0360, respectively). Since higher MFI scores indicate more severe fatigue, these associations would suggest that higher EV levels accompany a reduced fatigue burden in this group.

None of the EV markers reached nominal significance with symptoms in the COD and LC-19 groups (**Figures 5C and 5D**) (see **Supplementary Table S13** for details).

#### EV marker-blood parameter correlations

With respect to correlations between EV marker levels and clinical blood parameters, we should highlight that in the ME/CFS group (**Figure 6A**), all markers were negatively correlated with creatine phosphokinase (CPK) [CX3CR1 (R = −0.758, p = 0.0111); CD74 (R = −0.721, p = 0.0186); CD63 (R = −0.697, p = 0.0251), a relationship that has already been reported in the disease by our and other groups in larger cohorts (54,55);. Additional negative correlations of ACE1(CD143) and CX3CR1 with beta globulins were also identified in the ME/CFS group, indicating that increased levels of CD9/ACE1 and CD9/CX3CR1 EVs coincide with lower levels of beta-globulins which could indicate problems of the iron transport or the complement systems. A positive correlation of CD63 with ALT/GPT in these patients indicate liver function differences in patients with increased CD9/CD63 EV levels

In the FM group (**Figure 6B**), consistent correlations concentrated on renal and hematological parameters for all EV markers assessed. EV markers negatively correlated with creatinine [CD74 and CX3CR1 (both R = −0.854, p = 0.0017); ACE1(CD143) (R = −0.823, p = 0.0034); CD41/CD61 (R = −0.774, p = 0.0085); CD63 (R = −0.732, p = 0.0162); and CD13 (R = −0.701, p = 0.0239)], and positively correlated glomerular filtration rate (GFR), This indicates that FM patients with increased CD9 EVs may associate with muscle frailty or deconditioning, a status previously reported in these patients (61). Most EV markers were also positively correlated with red blood cell count, observations that may help stratify patients in RBC count-related fatigue.

In the COD group (Figure 6C), similar to ME/CFS all EV markers were strikingly negatively correlated with beta globulins [ACE1(CD143) (R = −0.821, p = 0.0036); CD63 (R = −0.815, p = 0.0041); CX3CR1 (R = −0.802, p = 0.0052); CD13 and CD41/CD61 (both R = −0.790, p = 0.0065); and CD74 (R = −0.772, p = 0.0089)], with additional negative correlations for total proteins and ferritin. This suggests that COD patients with increased CD9 levels hold marked alterations in iron transport or complement-related pathways.

In the LC-19 group (Figure 6D), four EV markers appeared negatively correlated with platelet count [CD13 (R = −0.891, p = 0.0005); CX3CR1 and ACE1(CD143) (R = −0.855, p = 0.0016); and CD74 (R = −0.818, p = 0.0038)], an observation that could help stratify patients with potential coagulation defects. Top discriminant single CD41/CD61 EV marker for this LC-19 group specifically shows a strong positive erythrocyte sedimentation rate (ESR) (R = 0.661, p = 0.0376), a nonspecific marker of systemic inflammation that could plausibly explain part of the symptom burden in this group. These findings may facilitate the identification of biologically distinct disease endotypes and help stratify patients with anemia-related or inflammatory processes.

## DISCUSSION

To our knowledge, this is the first study that evaluates the relative abundance of EV immune and circulatory markers coexisting with the exosomal CD9 tetraspanin and their potential correlations with patient symptoms and clinical blood parameters in a finely phenotyped cohort of patients that had received the clinical diagnosis of three diseases (ME/CFS, FM and LC-19). Differential diagnosis of these diseases can be confounded in the clinic by clinicians unfamiliar with these disorders because of symptom overlaps. In addition, the co-diagnosis of ME/CFS and FM is quite prevalent, and patients having received the diagnosis of LC-19 can fulfill ME/CFS clinical criteria within a few months post-infection (5,6), further complicating the task. Given that EVs can serve as sensors of the status of the organism, our aim was to determine whether our battery of selected EV markers could provide a basis for such differential diagnostic method and/or patient stratification to facilitate the diagnosis process in the clinic and as a guide towards the development of specific treatments. Despite the pilot nature of the study, it interestingly identifies distinct EV signatures across the four study groups (ME/CFS, FM, COD, and LC-19), revealing coordinated alterations that converge on a dysregulated platelet-immune-vascular axis. By integrating EV abundance, size distribution, and targeted surface marker profiling, our findings provide a system-level characterization of disease-associated processes rather than isolated biomarker changes.

EVs are increasingly recognized as functional mediators of intercellular communication, and their composition reflects the physiological state of their cells of origin (10,32,33). In this context, the EV profiles described here likely capture integrated signals from multiple cellular compartments affected in these patients, offering a multidimensional readout of systemic dysregulation. Importantly, this integrative approach reveals that EV-associated markers do not act as independent indicators but rather reflect coordinated changes across interconnected biological pathways. Previous studies have demonstrated EV differences in ME/CFS highlighting their importance to enlighten the pathomechanisms of the disease (11,13).

A key strength of this study lies in the direct comparison of clinically overlapping conditions (5,7,34). While the ME/CFS group showed partial or selective alterations in EV-associated markers, the LC-19 group exhibited the most pronounced and coordinated signature across the multiple pathways examined. This distinction suggests that LC-19 may represent a state of sustained multisystem activation, characterized by concurrent immune, vascular, and thrombotic dysregulation (6).

The increased abundance of CD9/CD41/CD61-positive EVs in both LC-19 and ME/CFS represents one of the most robust findings of this study and suggests the presence of shared pathophysiological mechanisms. These platelet-associated vesicles are indicative of enhanced platelet activation and are known to support pro-thrombotic and pro-inflammatory processes (35), consistent with accumulating evidence of microvascular dysfunction and microthrombotic events in both conditions (36). Beyond serving as biomarkers of platelet activation, EVs may actively contribute to thrombo-inflammatory signaling, further supporting their potential involvement in disease pathogenesis.

The concordant elevation of platelet-derived EVs in LC-19 and ME/CFS aligns with their classification as post-viral or post-infectious disorders and may reflect common biological consequences of infection-driven dysregulation (1,5,6,36,37). In contrast, the more heterogeneous EV profiles observed in FM and COD suggest greater mechanistic diversity. While FM is not traditionally considered a post-infectious condition, infection-triggered onset has been reported in a subset of patients, and the COD group may represent an overlapping phenotype that combines characteristics of both disorders.

An alternative interpretation is that the EV signature identified in LC-19 and ME/CFS reflects different stages of a shared pathological continuum. Under this hypothesis, LC-19 may represent a relatively earlier phase of post-infectious dysregulation, whereas ME/CFS corresponds to a more chronic state. This possibility is consistent with the longer disease duration of the ME/CFS cohort in the present study. Although speculative, such a temporal framework could help explain both the overlapping and distinct EV alterations observed across groups and should be explored in larger longitudinal studies.

On another side, immune-associated EV subpopulations were selectively enriched, particularly in LC-19. Importantly, although CD9/CD74- and CD9/CX3CR1-positive EVs are both linked to immune-related processes, they represent distinct functional pathways. CD74 is associated with antigen presentation and macrophage migration inhibitory factor (MIF)-mediated inflammatory signaling, reflecting sustained immune activation (38,39). In contrast, CX3CR1 is involved in leukocyte adhesion, migration, and vascular surveillance, particularly in non-classical monocytes (40–42). The concurrent increase of both EV populations in LC-19 therefore suggests a coordinated activation of inflammatory signaling and immune cell trafficking, pointing to a dynamic and spatially distributed immune response rather than a uniform inflammatory state.

Further supporting vascular involvement, EVs carrying components of the renin-angiotensin system showed disease-specific alterations. The observed increase in CD9/ACE1 (or CD143)-positive EVs in LC-19 (43–45), suggests a relative shift toward the pro-inflammatory and vasoconstrictive arm of the renin-angiotensin system. Given the central role of ACE2 in SARS-CoV-2 infection and vascular homeostasis, this imbalance could contribute to endothelial dysfunction and persistent vascular stress in LC-19 (46). Unexpectedly, the levels of ACE2 in LC-19 samples presenting increased ACE1 levels appeared unaltered. Although compensatory effects cannot be ruled out, the finding seems to agree with a recent report communicating no significant differences in soluble ACE/ACE2/angiotensin II between LC-positive and LC-negative individuals, noting that circulating levels may not reflect local tissue RAS activity (47). EV-bound ACE1 likely captures a distinct, tissue-proximal compartment not detectable by bulk soluble assays, which may explain the discrepancy. These findings reinforce the concept that EV-associated proteins can reflect dysregulation of regulatory systems involved in disease pathophysiology.

Although CD63 was included in the panel as a general tetraspanin marker of EV identity, it is also a well-established marker of platelet activation: CD63/LAMP-3 resides in the membrane of platelet dense granules and lysosomes and is translocated to the platelet surface specifically upon degranulation, being nearly undetectable on resting platelets but strongly upregulated following stimulation (48,49). Thus, the higher CD9/CD63-positive EV trend observed in ME/CFS and LC-19 may offer converging, independent support for platelet activation in these patient groups, complementing the CD41/CD61 findings. However, as CD63 is not platelet-restricted, future co-positivity analysis (e.g., CD63/CD41 or CD63/CD61 double-positive EV populations) would be required to confirm a platelet-derived origin rather than a broader shift in total small EV output (50,51).

Taken together, the combined alterations observed across platelet-derived, immune-associated, and RAS-related EV populations support the existence of an interconnected platelet-immune-vascular axis in these patients. Rather than independent processes, these pathways appear to be activated simultaneously, suggesting a coordinated network of dysregulation underlying disease pathology. EV profiling uniquely enables the dissection of this network into functionally distinct components, revealing pathway-specific alterations that would not be captured by bulk measurements.

Importantly, the differential EV signatures observed between disease groups highlight their potential clinical relevance. While some overlap exists between ME/CFS and LC-19 groups, LC-19 group displays a more distinct EV profile, particularly in immune- and vascular-associated markers. In this context, ROC analysis supports the potential utility of EV-based signatures for patient stratification and differentiation of clinically overlapping conditions. However, these signatures should be considered complementary to existing clinical frameworks rather than standalone diagnostic tools.

With respect to EV marker correlation with clinical variables and questionnaire-based patient phenotyping, we highlight two features. First, the significant associations did not occur randomly across the matrix but instead concentrated on one or two clinical variables within each group, with almost the entire marker panel tracking in the same direction. Second, the variables involved differed between groups: SF-36 Role Physical and CPK in ME/CFS (54,55); MFI Reduced Activity together with creatinine and glomerular filtration rate in FM; beta globulins in COD; and platelet count in LC-19. This convergence of all EV markers on a single variable remains more consistent with the panel behaving as a single shared axis of the EV surface signal than with independently regulated pathways.

Negative correlations between EV marker levels and platelet count in the LC-19 group (CD13, ρ = −0.891) constituted the strongest single association of the analysis. Residual platelet carryover into platelet-poor plasma would produce results in the opposite direction, and, as such, provide evidence against the presence of a simple pre-analytical artifact. Moreover, the platelet-specific marker CD41/CD61 representing the only panel member to show no association with platelet count in this group, argues against a straightforward platelet-consumption interpretation in which circulating platelets become depleted as platelet-derived EVs accumulate. Therefore, the association can best be described as between overall EV surface marker signal and circulating platelet count.

Several of the associated clinical parameters are not fully independent. For example, eGFR is calculated from serum creatinine, while beta-globulins and total protein levels are biologically related because beta-globulins constitute a fraction of total serum proteins. Ferritin is also linked to these parameters through shared influences such as inflammatory and metabolic status.

Of note, the scarcity of symptom-specific correlations does not weaken the abundance findings. Biomarkers that separate disease from healthy states do not always track severity linearly, a pattern also documented in Alzheimer’s plasma biomarkers (52). In addition, with 10 patients per group, the analysis only had the power to detect very large monotonic effects. Therefore, we may have failed to detect moderate but real associations and should not view their absence as evidence against their potential influence.

Plasma EV surface marker profiling to separate B-cell, monocyte/macrophage and NK subsets has recently been reported in LC-19 and ME/CFS (53); to our knowledge, however, this is the first study to profile EV-associated markers for platelet, immune, and renin-angiotensin systems simultaneously across four clinically overlapping conditions, and to identify both shared and disease-specific signatures.

Future studies should validate these findings in larger, more diverse, cohorts, including longitudinal analyses to assess temporal dynamics. Functional characterization of EVs will also be essential to determine their role as active mediators of disease, beyond their value as biomarkers.

Importantly, EV profiling enabled the identification of disease-specific patterns that may aid in distinguishing conditions with similar clinical presentations. While further validation is required, these findings underscore the potential of EV-based approaches for patient stratification and improved understanding of these chronic multisystem disorders.

## ACKNOWLEDGEMENTS

We want to particularly acknowledge the patients, and the Biobank IBSP-CV integrated in the ISCIII Biobanks and Biomodels Platform and in the Valencian Biobanking Network for their collaboration. MJV would like to thank Dr. Stuart P. Atkinson for reviewing the manuscript. This work was supported by the UCV predoctoral fellowship (VIDI) to ML; the Generalitat Valenciana CIAICO, grant number 2021/103 and the Spanish Ministry of Science and Innovation (PID2023-153299OB-I00) to EO; and the H2020 EU-OpenScreen-DRIVE project (grant agreement No. 823893) to MJV. Our screening platform forms part of the European research infrastructure ERIC EU-OpenScreen and of the national network TeNTaCLES (RED2018-102411-T). Part of the equipment employed was funded by the Generalitat Valenciana and co-financed with FEDER funds (PO FEDER of Comunitat Valenciana 2014–2020).

